# Xenogeneic regulation of the ClpCP protease of *Bacillus subtilis* by a phage-encoded adaptor-like protein

**DOI:** 10.1101/569657

**Authors:** Nancy Mulvenna, Ingo Hantke, Lynn Burchell, Sophie Nicod, Kürşad Turgay, Sivaramesh Wigneshweraraj

**Affiliations:** MRC Centre for Molecular Bacteriology and Infection, Imperial College London, London, SW7 2AZ, UK; Institute für Mikrobiologie, Leibniz Universität Hannover, Herrenhäuser Str. 2, 30419 Hannover, Germany

**Author notes:** **Corresponding author:** Sivaramesh Wigneshweraraj, MRC Centre for Molecular Bacteriology and Infection, Imperial College London, London SW7 2AZ, UK.

**Keywords:** *Bacillus subtilis*, SPO1 phage, ClpCP protease, proteolysis, host takeover, bacterial growth

## Abstract

SPO1 phage infection of *Bacillus subtilis* results in a comprehensive remodelling of processes leading to conversion of the bacterial cell into a factory for phage progeny production. A cluster of 26 genes in the SPO1 genome, called the host takeover module, encodes for potentially cytotoxic proteins for the specific shut down of various host processes including transcription, DNA synthesis and cell division. However, the properties and bacterial targets of many genes of the SPO1 host takeover module remain elusive. Through a systematic analysis of gene products encoded by the SPO1 host takeover module we identified eight gene products which attenuated *B. subtilis* growth. Out of the eight gene products that attenuated bacterial growth, a 25 kDa protein, called Gp53, was shown to interact with the AAA+ chaperone protein ClpC of the ClpCP protease of *B. subtilis*. Results reveal that Gp53 functions like a phage encoded adaptor protein and thereby appears to alter the substrate specificity of the ClpCP protease to modulate the proteome of the infected cell to benefit efficient SPO1 phage progeny development. It seems that Gp53 represents a novel strategy used by phages to acquire their bacterial prey.

**Significance statement:** Viruses of bacteria (phages) represent the most abundant living entities on the planet, and many aspects of our fundamental knowledge of phage–bacteria relationships remain elusive. Many phages encode specialised small proteins, which modulate essential physiological processes in bacteria in order to convert the bacterial cell into a ‘factory’ for phage progeny production – ultimately leading to the demise of the bacterial cell. We describe the identification of several antibacterial proteins produced by a prototypical phage that infects *Bacillus subtilis* and describe how one such protein subverts the protein control system of its host to benefit phage progeny development. The results have broad implications for our understanding of phage–bacteria relationships and the therapeutic application of phages and their gene products.

---

Much like eukaryotic and archaeal viruses, which derail the host’s cellular processes to facilitate viral replication, phages have evolved complex strategies to acquire their bacterial hosts. In order to successfully infect and replicate in the bacterial cell, many phages encode proteins that specifically interfere with essential biological processes of the host bacterium, including transcription, translation, DNA replication and cell division (1). Phage proteins that interfere with host processes are typically small in size (on average ∼160 amino acid residues) and are usually produced at high levels early in the infection cycle (2). SPO1 is a prototypical lytic phage of *Bacillus subtilis* and its genes are categorised as early, middle and late to reflect the time of their expression during SPO1 development in *B. subtilis*. The majority of SPO1 early genes associated with host takeover are in the 12.4 kb terminal region of the genome, which includes the 26-gene host takeover module (Fig. 1*A*) (3, 4). The genes within the host takeover module, *gp37-gp60*, have several hallmarks to suit the characteristics of phage proteins that interfere with host processes: They are mostly small, produced early in infection and contain promoters and ribosome binding sites characteristic of highly expressed genes (3, 5). Many of them have been previously shown to be involved in the shut-off of bacterial DNA and RNA synthesis (*gp38*, *gp39*, *gp40*, *gp44*, *gp50* and *gp51*) or to inhibit cell division (*gp56*) during SPO1 infection (6–8). Further, plasmid-borne expression of *gp44* and *gp56* in *B. subtilis* has been shown to attenuate growth and reduce viability, respectively (8, 9). With the exception of the product of *gp44*, which has been postulated to interact with *B. subtilis* RNA polymerase (9, 10), the bacterial targets and mechanism of action of the gene products encoded by the host takeover module of SPO1 remain elusive. Clearly, phages and their gene products represent an underexploited resource for potentially developing novel antibacterial strategies and to gain new insights into bacterial cell function and regulation. In this study, we undertook a systematic approach to identify genes in the SPO1 phage host takeover module that had a detrimental effect on *B. subtilis* growth and unveil the biological role of the product of *gp53*, which interacts with the Hsp100/Clp family member ClpC of *B. subtilis*.

**Fig. 1.**
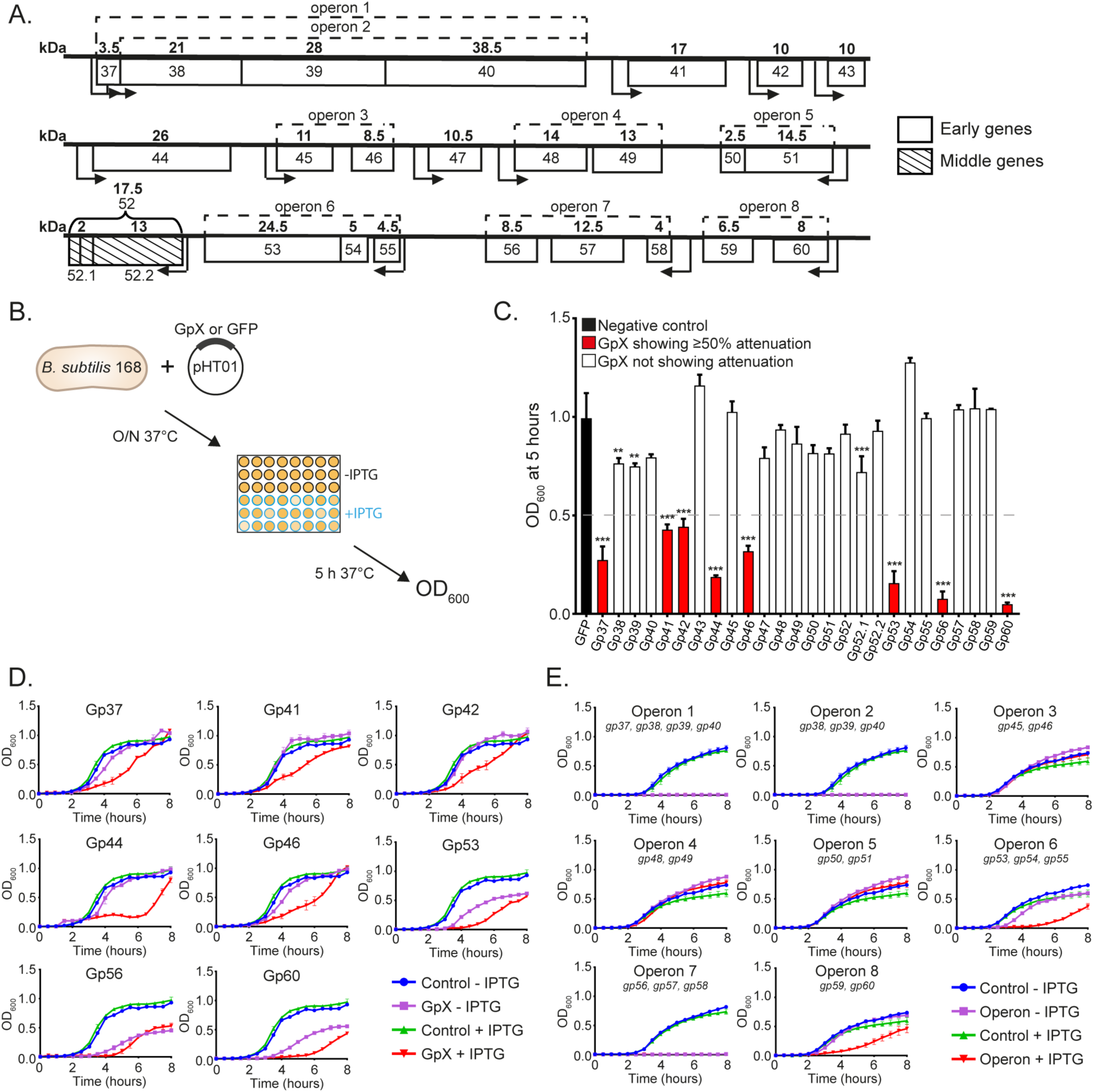
SPO1 host takeover module genes that attenuate *B. subtilis* growth. (*A*). Schematic of the SPO1 host takeover module. The molecular weights (kDa) of the individual gene products are shown above each gene in bold and operons are indicated by dotted lines. The predicted positions of promoters are shown as arrows indicating the direction of transcription. (*B*). Schematic of the experimental procedure used to identify SPO1 host takeover module gene products that attenuate the growth of *B. subtilis*. (*C*). Graph showing the OD_600_ values of *B. subtilis* cultures at 5 hours of growth in the presence of IPTG which induces the expression of the individual host takeover module genes. Gene products shown in red displayed ≥50% attenuation when compared to control cells expressing GFP. (*D*). Graphs showing growth curves (in red) of *B. subtilis* cultures expressing SPO1 host takeover module genes that attenuated growth ≥50%; control growth curves are indicated in the key. (*E*). Graphs showing growth curves (in red) of *B. subtilis* cultures expressing the individual operons of the SPO1 host takeover module; control growth curves are indicated in the key. Error bars in *C*, *D* and *E* represent SEM (n=3). Statistical analyses were performed by one-way ANOVA (** P<0.01, *** P<0.001).

## Results

### The effect of SPO1 host takeover module genes on *B. subtilis* growth

We wanted to identify genes in the SPO1 host takeover module that had a detrimental effect on *B. subtilis* growth by growing bacteria in the absence and presence of isopropyl β-D-1- thiogalactopyranoside (IPTG), which allowed plasmid pHT01 (11) borne expression of the 26 host takeover genes either individually or with other genes in their respective operons (Fig. 1*A*). Any effect of the gene products of the host takeover module on *B. subtilis* growth was monitored by determining the cell density by measuring light absorbance of the culture at 600 nm after a 5-hour period of incubation at 37°C (Fig. 1*B*). As the control, we used bacteria containing pHT01 plasmid expressing green fluorescent protein (GFP). As shown in Fig. 1*C*, when the SPO1 phage host takeover module genes were expressed individually in *B. subtilis*, the growth of bacteria expressing Gp37, Gp41, Gp42, Gp44, Gp46, Gp53, Gp56 and Gp60 was attenuated by 50% or more when compared to the control cells expressing GFP. The individual graphs in Fig. 1*D* show growth curves of *B. subtilis* expressing Gp37, Gp41, Gp42, Gp44, Gp46, Gp53, Gp56 and Gp60 over a period of 8 hours. We noted that, under our conditions, the plasmid-borne expression of Gp37, Gp41, Gp42, Gp44, Gp46, Gp53, Gp56 and Gp60 did not inhibit growth *per se* but attenuated growth by extending the lag time to varying degrees (Fig. 1*D*). Further, it seemed that leaky expression (which occurs in the absence of the inducer) of Gp53, Gp56 and Gp60 also attenuated growth to some degree, indicating that the latter SPO1 gene products are potentially more toxic to *B. subtilis* than the others (i.e. Gp37, Gp41, Gp42, Gp44 and Gp46). The expression of the SPO1 host takeover module genes together with other genes in their respective operons revealed that operons containing genes shown to attenuate growth when expressed individually also attenuated growth efficiently (Fig. 1*E*) with the following exceptions: Firstly, Gp38, Gp39 and Gp40 when expressed together in operon 1 and operon 2, appeared to act synergistically and displayed an enhanced ability to attenuate bacterial growth (compare Fig. 1*C* and Fig. 1*E*). Secondly, we note that in *B. subtilis* cells in which the host takeover module genes in operon 1, 2 and 7 are expressed together do not recover under our experimental conditions. This indicates that the host takeover module gene products within each operon functionally interact and thus have a more pronounced effect on host physiology than when expressed individually. Finally, we note that Gp46 is no longer able to attenuate growth of *B. subtilis* when expressed together with Gp45 in operon 3. This implies that the Gp45 somehow mitigates the antagonistic effect of Gp46 on *B. subtilis* cells. Overall, we conclude that recombinant forms of Gp37, Gp41, Gp42, Gp44, Gp46, Gp53, Gp56 and Gp60 have a detrimental effect on *B. subtilis* growth in the absence of SPO1 infection, presumably by targeting essential cellular processes.

### Gp53 interacts with the ClpC ATPase of the ClpCP protease in *B. subtilis*

Since Gp53 was experimentally better tractable than the other SPO1 host takeover module gene products, we focused on identifying the target(s) of Gp53 in *B. subtilis*. We constructed an amino (N) terminal hexa-histidine (6His) tagged version of Gp53 to identify its bacterial target(s) by conducting a pull-down assay using whole-cell extracts of exponentially growing *B. subtilis* cells. Initially, we investigated whether the histidine tagged version of Gp53 retained its ability to attenuate *B. subtilis* growth under the conditions described in Fig. 1. As shown in Fig. 2*A*, the activity of N terminal 6His tagged Gp53 and its untagged counterpart did not differ significantly. For simplicity, from here on the N terminal 6His tagged version of Gp53 will be referred to as Gp53. In order to perform the pull-down assays, purified Gp53 was immobilised onto nickel resin and the ‘charged’ resin was incubated with whole-cell extracts prepared from exponentially growing *B. subtilis* cells (Fig. 2*B*). The resin was then extensively washed to remove any non-specific interactions before analysis by SDS polyacrylamide gel electrophoresis (PAGE). As shown in Fig. 2*C*, when the pull-down assay was conducted in the presence of Gp53, we detected a specific enrichment of a band on the SDS-PAGE gel (Fig. 2*C*, arrow in lane 3), which was not observed in the control reactions with ‘uncharged’ resin (i.e. in the absence of any immobilised protein) (Fig. 2*C*, lane 2). The enriched band was investigated by linear quadrupole ion trap Fourier transform mass spectrometry (LTQ-FTMS) analysis, which revealed it to be the Hsp100/Clp family member ClpC, the ATPase subunit of the ClpCP protease in *B. subtilis*. To further validate that Gp53 interacts with ClpC, we repeated the pull-down assay using purified carboxyl terminal FLAG tagged ClpC and nickel resin with immobilised Gp53. As shown in Fig. 2*D*, FLAG tagged ClpC appears to weakly interact with the nickel resin (lane 4) in the absence of Gp53. However, a specific enrichment of ClpC is clearly seen in the presence of Gp53 (Fig. 2*D*, lane 3).

**Fig. 2.**
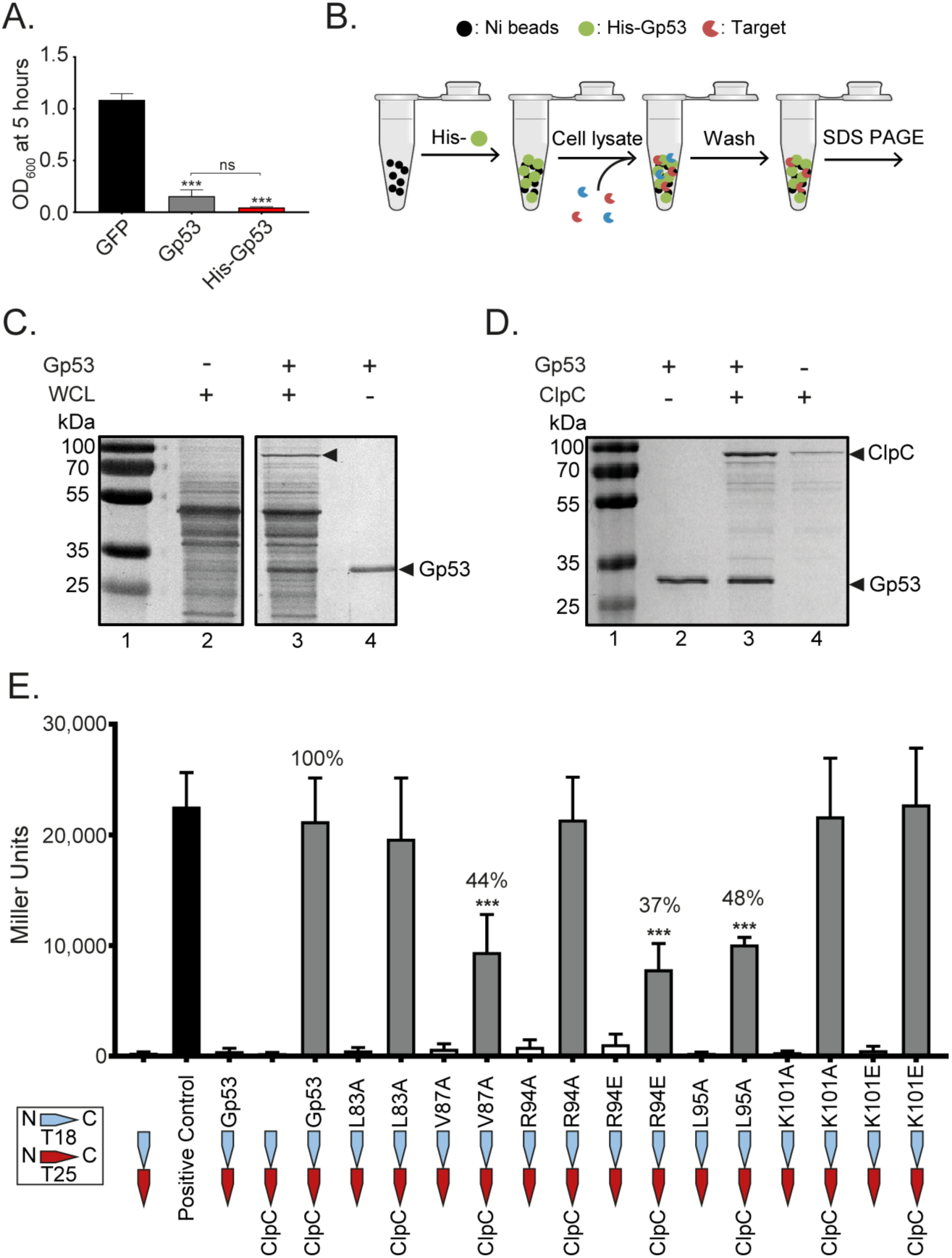
Gp53 interacts with the ClpC ATPase of the ClpCP protease in *B. subtilis*. (*A*). Bar chart comparing the efficacy of growth attenuation of a culture of *B. subtilis* either expressing N terminal 6His tagged Gp53 (red) or untagged Gp53 (grey). (*B*). Schematic of the pull-down assay used to identify the bacterial target(s) of Gp53. (*C*). A representative image of a SDS-PAGE gel showing results of the pull-down assay with Gp53 and whole-cell extracts (WCL) of *B. subtilis*. The band specifically enriched in reactions containing immobilised Gp53 is indicated with an arrow in lane 3. (*D*). A representative image of a SDS-PAGE gel showing results of pull down assay with purified Gp53 and N terminal FLAG tagged ClpC. The migration positions of Gp53 and ClpC are indicated. E. Bar chart showing the results from the bacterial two-hybrid interaction assay with ClpC and mutant variants of Gp53. The ClpC binding activity of the Gp53 mutants is shown as a percentage of wild-type Gp53 activity above the bars of mutants with compromised binding activity. Error bars in *A* and *E* represent SEM (n=3). Statistical analyses were performed by one-way ANOVA (ns not significant, *** P<0.001).

To establish that the interaction between Gp53 and ClpC is specific and to identify amino acids in Gp53 important for binding to ClpC, we conducted a BLAST search using standard search parameters and SPO1 Gp53 as a query sequence. Three homologous proteins and 1 protein fragment from SPO1 related phages were found (Fig. S1) with amino acids (L83, V87, R94, L95 and K101) conserved across all five sequences. All these residues were individually substituted with alanine (A), apart from the positively charged residues R94 and K101, which were also replaced with negatively charged glutamic acid (E) residues. Next, a bacterial two-hybrid (BTH) interaction assay was performed to determine how the amino acid substitutions in Gp53 affected its ability to interact with ClpC. We opted for the bacterial adenylate cyclase two-hybrid (BACTH) system in which both genes *gp53* and *clpC* were co-expressed in a Δ*cya E. coli* strain DHM1 as fusions to one of two fragments (T18 and T25) of the catalytic domain of *Bordetella pertussis* adenylate cyclase (12). Interaction of two-hybrid proteins results in a functional complementation between T18 and T25 leading to cAMP synthesis, and transcriptional activation of the lactose operon that can be detected in a β-galactosidase assay. As shown in Fig. 2*E*, reactions with Gp53 variants harbouring an alanine substation at V87A or L95A and charge-reversal substitution at R94 (R94E) displayed significantly lower β-galactosidase activity compared to the reaction with wild-type Gp53. We conclude that proximally located amino acid residues V87, R94 and L95 in Gp53 are important determinants for binding to ClpC.

### Gp53 stimulates the ATPase activity of ClpC in an analogous manner to *B. subtilis* adaptor proteins

The role of ClpC in *B. subtilis* is the ATP hydrolysis dependent unfolding and loading of substrate proteins for degradation by the protease ClpP. Substrate specificity upon ClpC is conferred by different adaptor proteins, which interact with ClpC and cause ClpC to oligomerise and form a complex with ClpP monomers that come together to form the proteolytic chamber (Fig. 3*A*). In other words, the adaptor protein is an obligatory activator of the ClpCP protease (13). Since the binding of the adaptor protein, such as the well-documented MecA protein, has been shown to stimulate the basal ATPase activity of ClpC, we initially tested how Gp53 binding affected the ATPase activity of ClpC. Results shown in Fig. 3*B* indicated a dose dependent stimulation of the ATPase activity of ClpC by Gp53. Control experiments with the mutant variant of Gp53 harbouring the R94E substitution, which displayed a compromised ability to bind ClpC in the BTH assay (Fig. 2*E*), revealed that the stimulation of ClpC’s basal ATPase activity was due to the specific action of Gp53 (Fig. 3*C*). We next wanted to determine if Gp53 competed with native adaptor proteins for binding to ClpC. Using MecA as a model adaptor protein (14–16), we initially conducted ATPase assays to determine whether Gp53 and MecA can bind simultaneously to ClpC and can act synergistically to stimulate the basal ATPase activity of ClpC. As shown in Fig. 3*D*, the addition of MecA (reaction I) or Gp53 (reaction II) resulted in the stimulation of the basal ATPase activity of ClpC. However, the presence of MecA and Gp53 together in the reaction, regardless of the order of addition, did not result in an increase in ClpC’s ATPase activity to a level higher than the ATPase activity seen when MecA and Gp53 were added individually (Fig. 3*D*, compare reactions I and II with III and IV). Therefore, we conclude that, although MecA and Gp53 individually stimulate the basal ATPase activity of ClpC, they compete for binding to ClpC with comparable efficacy.

**Fig. 3.**
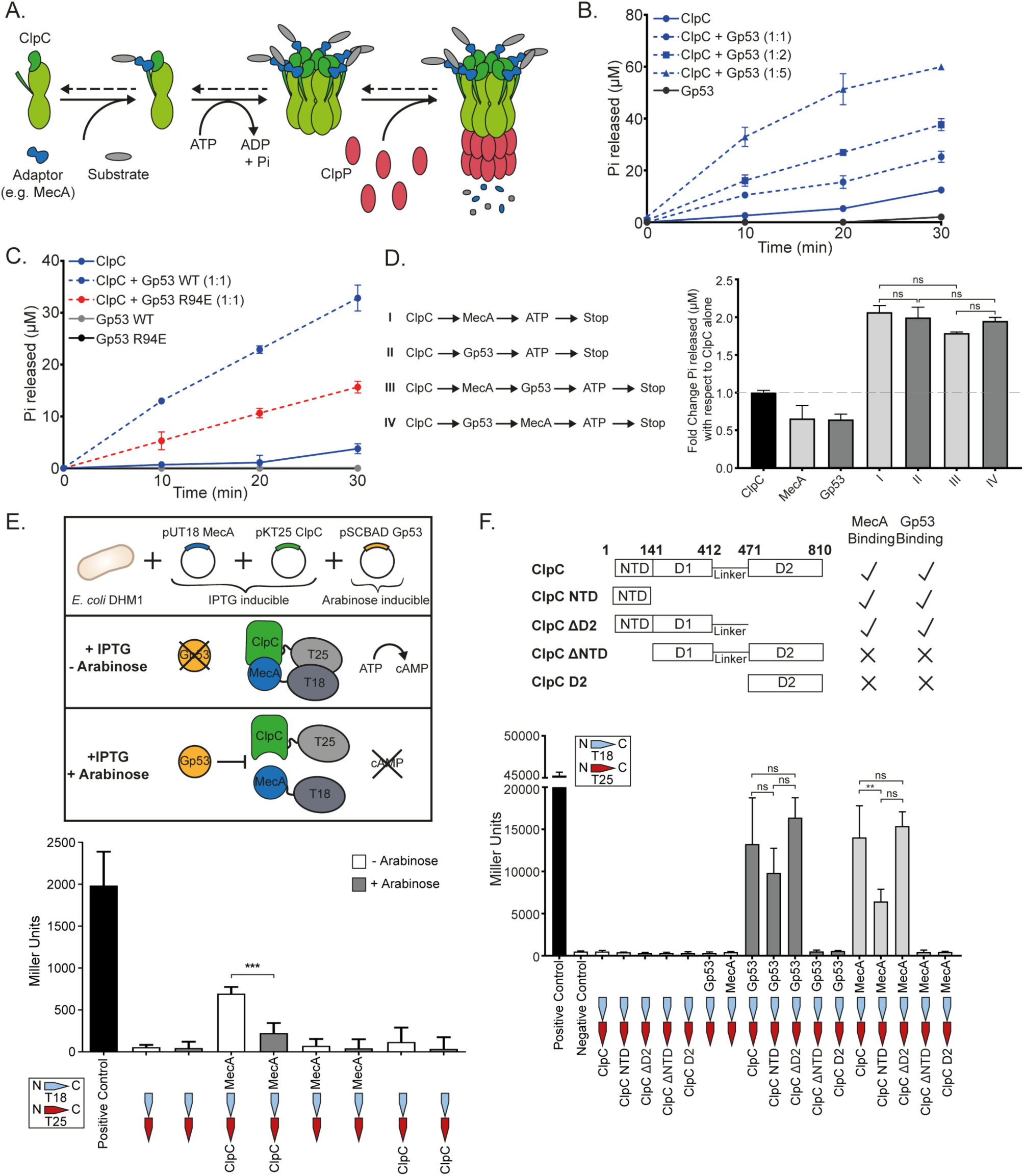
Gp53 stimulates the ATPase activity of ClpC and competes with *B. subtilis* adaptor protein MecA for binding to ClpC. (*A*). Schematic showing how the ATP hydrolysis and adaptor protein mediated formation of the functional ClpCP protease in *B. subtilis*. Adapted from Molière et al. (28) (*B*). Graph showing the amount of ATP hydrolysed (phosphate (Pi) release in µM) as a function of time by ClpC (0.2 µM) alone and in the presence of different amounts of Gp53 (0.2 µM, 0.4 µM, 1 µM). (*C*). As in *B*. but including Gp53 R94E (0.2 µM). (*D*). Bar chart showing results from the ATPase assay (as in Fig. 3B) in which ClpC (50 nM) was incubated with equimolar amounts of MecA (reaction I), Gp53 (reaction II) or MecA and Gp53 (added to the reaction in different orders; reactions III and IV). The amount of Pi released (µM) is expressed as fold change with respect to reaction with ClpC alone i.e. its basal ATPase activity. (*E*). Bar chart showing the results from the modified bacterial two-hybrid interaction assay to demonstrate that Gp53 competes with MecA for binding ClpC. The schematic on the top shows the assay setup (see text for details). (*F*). Bar chart showing the results from the bacterial two hybrid assay showing the binding of Gp53 or MecA to different domains of ClpC (as shown in the schematic at the top). In *B*-*F*, error bars represent SEM (n=3). Statistical analyses were performed by one-way ANOVA (ns not significant, ** P<0.01, *** P<0.001).

To directly determine that Gp53 competes with MecA for binding to ClpC, we used a modified version of the BTH assay described in Fig. 2*E*. In this assay, MecA and ClpC were fused to the T18 and T25 fragments, respectively, of the catalytic domain of *B. pertussis* adenylate cyclase and transformed into the Δ*cya E. coli* strain DHM1containing a plasmid in which Gp53 expression was under the control of the L-arabinose inducible *araB* promoter. We expected that if Gp53 competed with MecA for binding to ClpC, then the productive interaction between MecA and ClpC would be disrupted when the expression of Gp53 is induced with L-arabinose (Fig. 3*E*, see schematic). As expected, the results revealed that the β-galactosidase activity originating from the productive interaction between MecA and ClpC was reduced by ∼3 fold in the presence of L-arabinose (Fig. 3*E*). Further, previous studies (17–20) revealed the NTD domain (amino acid residues 1-141) and a linker region (amino acid residues 412-471) in ClpC to be important for binding to adaptor proteins MecA and McsB. Therefore, to determine whether the Gp53 and MecA/McsB interacting regions on ClpC overlap or are different, we fused four fragments of ClpC (Fig. 3*F*, see schematic) to T25 and used either MecA or Gp53 fused to T18 in the BTH assay. Results shown in Fig. 3*F* clearly reveal that both MecA (and by extension McsB; see below) and Gp53 bind to overlapping surfaces on ClpC with the linker region being of less importance for binding Gp53. In summary, we conclude that Gp53, although it clearly activates ClpC in an analogous manner to *B. subtilis* adaptor proteins, it is likely to compete with them for binding to ClpC (see below). Thus, by inference, we suggest that Gp53 could affect the normal functioning of the ClpCP protease by excluding the functionally obligatory adaptor proteins from interacting with it.

### Gp53 alters the specificity of the ClpCP protease in *B. subtilis*

We next investigated the effect of Gp53 on the protease activity of ClpCP. Although adaptor proteins like MecA are required for activation and to confer specificity upon the ClpCP protease, they are degraded along with the substrate or even in the *absence* of the substrate (15). Therefore, to determine whether Gp53 *inhibits* the proteolytic activity of the ClpCP protease or merely *alters* its specificity during SPO1 development in *B. subtilis*, we initially tested, using purified components, whether Gp53 is also degraded by the ClpCP protease in the absence of any substrate. As shown in Fig. 4*A*, in the absence of any substrate, MecA, as expected, was degraded by ClpCP protease (left panel). Similarly, Gp53 was also degraded, albeit at a slower rate than MecA, by the ClpCP protease (Fig. 4*B*; right panel). Further, consistent with the results in Fig. 3, results in Fig. 4*B* confirmed that both Gp53 and MecA compete for binding to ClpC, since the addition of both proteins together resulted in an overall decreased rate of degradation of either protein (compare lanes 3, 4, 6 and 7 from Fig. 4*A* with lanes 2 and 3 from Fig. 4*B*).

**Fig. 4.**
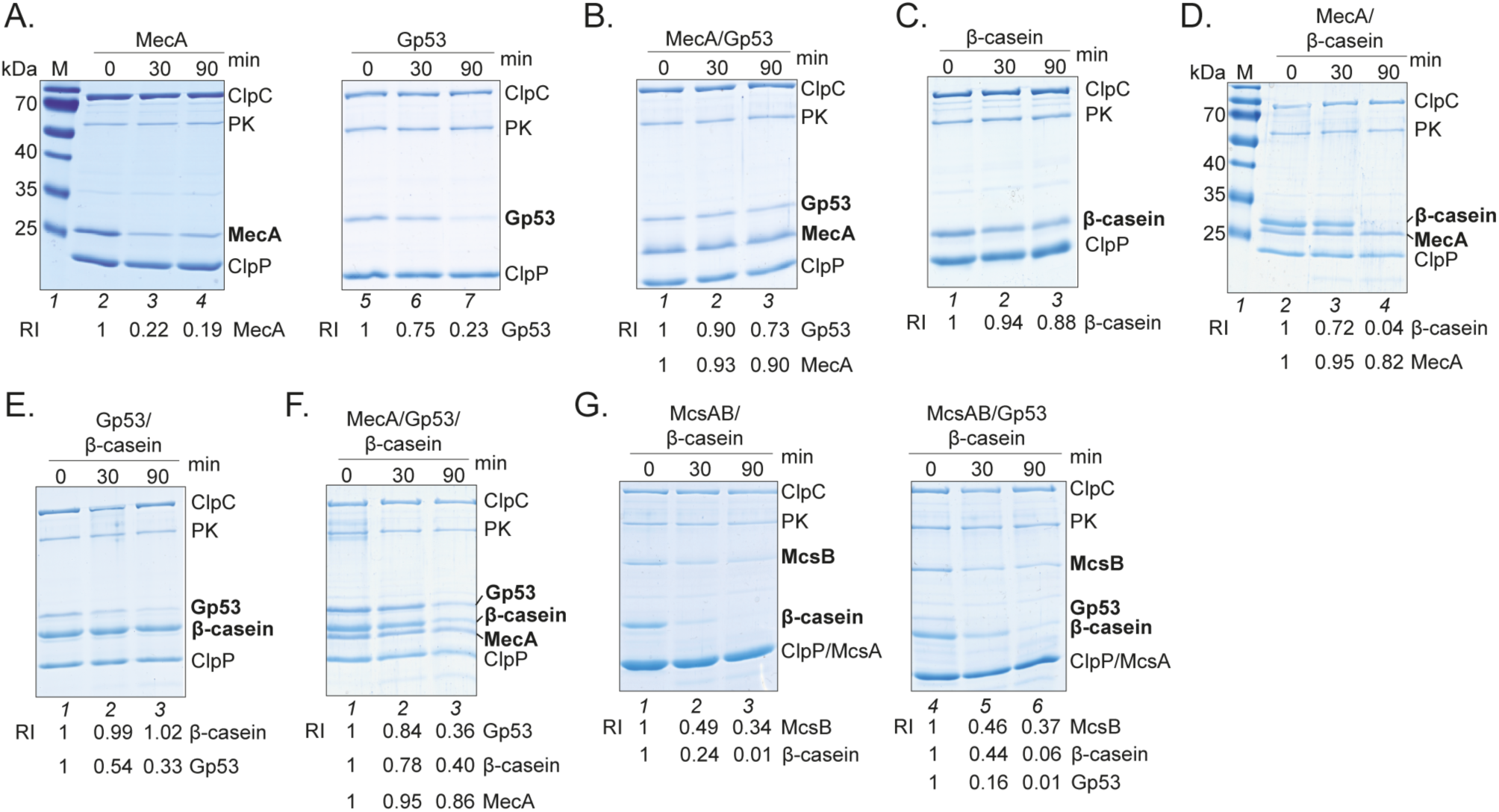
Gp53 alters the specificity of the ClpCP protease in *B. subtilis*. (*A*). Representative images of SDS-PAGE gels of *in vitro* degradation of MecA and Gp53 by ClpCP protease. Relative intensities (RI) of the bands corresponding to MecA or Gp53 are given below relative to the intensity of the ClpP band in the corresponding lanes. The migration position of ClpC (1 µM), MecA (1 µM), Gp53 (1 µM) and ClpP (1 µM) are indicated. Pyruvate kinase (20 ng/ml, PK, indicated) and phosphenolpyruvate (4 mM) were used as an ATP generation system. (*B*). As in *A*. but equimolar amounts of MecA and Gp53 were added together. (*C*). As in *A*. but the *in vitro* degradation assays were conducted in the presence of 3 µM β-casein and the absence of MecA or Gp53. (*D*). As in *C*. but the *in vitro* degradation assays were conducted in the presence of MecA. (*E*). As in *C*. but the *in vitro* degradation assays were conducted in the presence of Gp53. (*F*). As in *C*. but the *in vitro* degradation assays were conducted in the presence of MecA and Gp53. (*G*). As in *C*. but the in vitro degradation assays were conducted with McsA/B (1 µM each) in the absence and presence of Gp53.

Next, we conducted degradation assays using the intrinsically unfolded β-casein as a model substrate in the presence of MecA and/or Gp53. Consistent with previous studies, the control reaction in the absence of MecA or Gp53 did not result in the degradation of β-casein (Fig. 4*C*). However, degradation of β-casein was detected in the presence of MecA (compare lanes 1-3 from Fig. 4*C* with lanes 2-4 from Fig. 4*D*). Interestingly, although ClpC is activated by Gp53 (Fig. 3) leading to degradation of Gp53 by ClpCP (Fig. 4*A*), β-casein was not degraded in the presence of Gp53 (Fig 4*E*). This indicated that Gp53 is likely to alter the substrate specificity of ClpCP. In other words, Gp53 may not act as a regular adaptor protein but recognises other, yet unknown, substrates for proteolysis during SPO1 development in *B. subtilis*. Consistent with the results in Fig. 4*B*, the presence of Gp53 and MecA *together* in the reaction decreased the rate of β-casein degradation (Fig. 4*F*): following 90 minutes of incubation ∼10 fold β-casein was left intact compared to reactions without Gp53 (compare lane 4 in Fig. 4*D* with lane 3 in Fig. 4*F*). Additional experiments with a different adaptor protein, McsB/A (McsB requires McsA for activation (21)), confirmed that the competition for binding to ClpC by Gp53 was not restricted to MecA. As shown in Fig. 4*G*, the rate of degradation of β-casein in reactions with McsB/A was reduced in the presence of Gp53 when compared to reactions without Gp53 (compare lanes 4-6 with 1-3). In conclusion, the results unambiguously reveal that Gp53 competes with *B. subtilis* adaptor proteins for binding to ClpC and does not inhibit but is likely to alter the specificity of the ClpCP protease to benefit SPO1 development.

### Compromised ClpCP protease activity affects the efficacy of SPO1 development in *B. subtilis*

We posited that if the role of Gp53 is to alter the specificity of the ClpCP protease to allow successful development of SPO1 in *B. subtilis*, then a Δ*clpC B. subtilis* strain (IH25) would provide a compromised host environment for SPO1 development than wild-type *B. subtilis* cells. We compared the ability of SPO1 to lyse an exponentially growing culture of wild-type and Δ*clpC B. subtilis* by measuring cell density (light absorbance at OD_600_) as a function of time following SPO1 infection. The growth of wild-type and Δ*clpC* strains under our experimental conditions did not detectably differ (Fig. 5*A*). A rapid drop in cell density, indicating cell lysis, was observed after ∼30 minutes in the wild-type *B. subtilis* culture infected with SPO1 at OD_600_ 0.2 (Fig. 5*B*). As expected, the Δ*clpC B. subtilis* culture infected with SPO1 continued to grow for a further 20 minutes, reaching a higher cell density than the wild-type strain, before undergoing cell lysis (Fig. 5*B*). As shown in Fig. 5*C*, similar results were obtained with *B. subtilis* strains containing ClpC which is unable to hydrolyse ATP because of two mutations within the Walker B domain in both ATPase domains (strain IH140 (18)) or efficiently interact with ClpP because of a deletion in a region required for binding to ClpP (VGF::GGR, strain IH217 (22)): In the case of the IH140 and IH217 mutant *B. subtilis* strains, the culture continued to grow for a further 10 minutes compared to the wild-type culture before cell lysis occurred. Overall, the results are consistent with the findings above and indicate that the altering the specificity of ClpCP protease by Gp53, but not its inhibition, is required for optimal SPO1 development in *B. subtilis*.

**Fig. 5.**
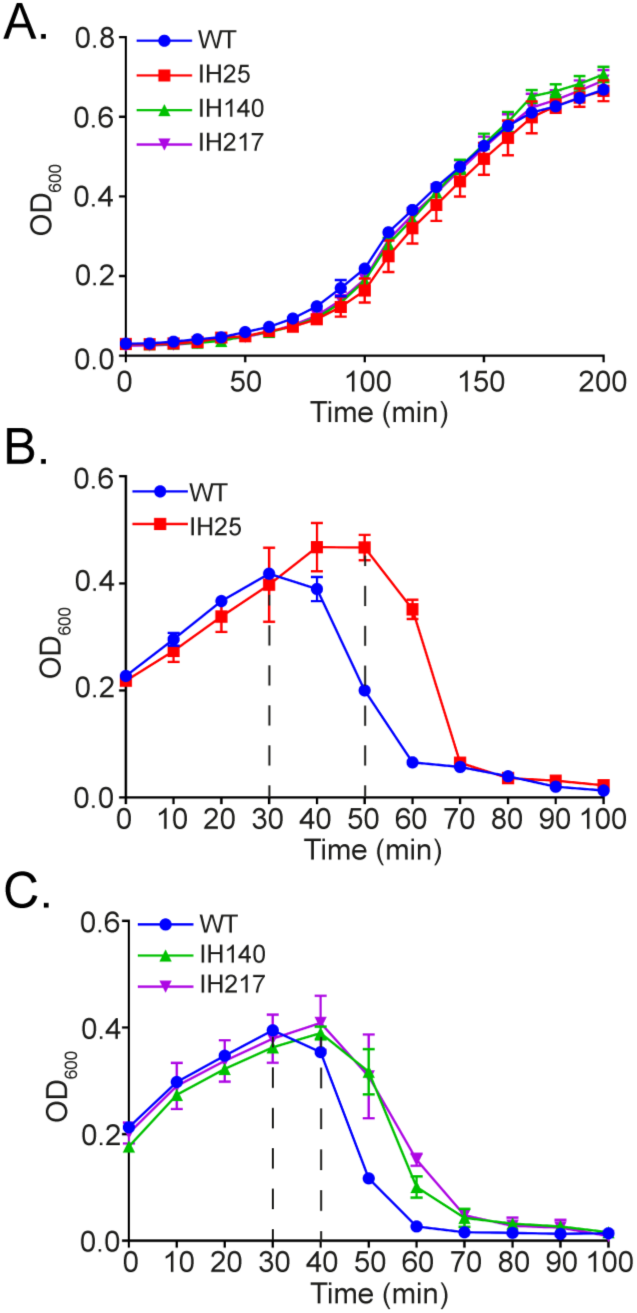
Compromised ClpCP protease activity affects the efficacy of SPO1 development in *B. subtilis.* (*A*). Graph showing the growth curves of wild-type (WT), IH25 (Δ*clpC*), IH140 (*clpC* DWB) and IH217 (*clpC* VGF::GGR) *B. subtilis* cultures. (*B*). Graph showing the optical density as a function of time of a culture of exponentially growing WT and IH25 *B. subtilis* cells following infection with SPO1 at OD_600_ 0.2. (*C*). As in *B*. but with WT, IH140 and IH217 *B. subtilis* cells. Error bars in *A*, *B* and *C* represent SEM (n=3).

## Discussion

A common theme by which phages affect host physiology to benefit phage progeny development is through the modulation or inhibition of bacterial cellular processes (1, 2). Previous studies (6–9) revealed that SPO1 infection results in the remodelling of several host processes by six (Gp38, Gp39, Gp40, Gp44, Gp50 and Gp51) of the twenty-six genes encoded by the host takeover module. Specifically, although the molecular details still remain elusive, Gp38, Gp39, Gp40, Gp44, Gp50 and Gp51 have been implicated in the shut-off of host macromolecular biosynthetic processes (RNA, DNA and protein synthesis) and Gp56 in the inhibition of bacterial cell division (6–8). This study revealed that Gp37, Gp41, Gp42, Gp44, Gp46, Gp53, Gp56 and Gp60 attenuate the growth of *B. subtilis* in the absence of SPO1 infection (Fig. 1). It seems that the individual effects of some host takeover module gene products, e.g. Gp38, Gp39, Gp40, might not be sufficient to affect bacterial growth. In support of this view, co-expression of Gp38, Gp39 and Gp40, which constitute operon 2 of the host takeover module (Fig. 1*A*), resulted in increased growth attenuation, presumably through synergistic activities of Gp38, Gp39 and Gp40. As phage genomes tend to be compact and efficient, it is remarkable that SPO1 has evolved many elaborate mechanisms to take over *B. subtilis* cells. We predict that the action of each individual host takeover module gene product is carefully regulated in a temporally coordinated manner and that some functionally interact with each other to bring about the desired effect (e.g. Gp38, Gp39 and Gp40) or control their functionalities. The observation that the co-expression of Gp45 with Gp46 (operon 3) counteracts the effect of the latter on *B. subtilis* growth (Fig. 1*E*) further underscores this view. Further, it is tempting to speculate that genes within operon 3 of the host takeover module are akin to a toxin/anti-toxin module. Further, it is important to remember that most studies on phage-host interactions, like the present one, are conducted under ‘optimal’ laboratory conditions. Thus, it is possible that some of the SPO1 host takeover module gene products might only be required for infecting and replicating in bacteria in different physiological states e.g. a nutrient starved population of bacteria. For example, Gray et al recently reported that *B. subtilis* can exist in an oligotrophic state without sporulating (23). It would thus be interesting to investigate whether some SPO1 host takeover gene product and their targets become essential for SPO1 development under this state of growth. Further, our earlier work on the T7 phage led to the identification of a T7 gene product involved in the inhibition of the bacterial RNAP only in the stationary phase of growth (24).

Although it is common for phages to *depend* on or *inhibit* the host’s protein degradation machinery for phage developmental requirements (e.g. lysis-lysogeny decision in phage lambda (25), DNA replication/transcription decision in phage Mu (26) or inhibition of Lon protease by T4 (27)), to the best of our knowledge, this study presents the only example of a phage protein that *alters* the substrate specificity of the host’s protein degradation machinery to allow optimal phage development. Under standard laboratory conditions, the absence of ClpCP protease activity had a subtle yet consistent detrimental effect on the efficacy of SPO1 development in *B. subtilis* (Fig. 5). Thus, it is possible that the requirement for the ClpCP protease activity by SPO1 becomes more prominent under more native and/or specific conditions for *B. subtilis* (see above). The results reveal that SPO1 Gp53 competes with host adaptor protein(s) for binding to ClpC and thereby alters the specificity of the ClpCP protease. Since different adaptor proteins, for example McsB and MecA, can compete for binding to ClpC to confer substrate specificity upon the ClpCP protease (14, 17) and it seems that Gp53 is an example of an adaptor-like protein produced by a phage. Consistent with this view, the results revealed that the binding site of Gp53 on ClpC is likely to overlap with that of native adaptor proteins such as MecA or McsB (Fig. 3 and Fig. 4). Thus, it is conceivable that Gp53 functionally mimics the role of a *B. subtilis* adaptor protein, which, consequently, could result in the subversion of the ClpCP protease to benefit phage development. We propose the following two mutually exclusive scenarios: (1) that Gp53 could act like an adaptor protein and target SPO1 derived substrates for proteolysis and consequently interferes with the recognition and targeting of “natural” substrates by the native (bacterial) adaptor proteins for proteolysis by the ClpCP protease and/or (2) Gp53 repurposes the ClpCP protease to degrade or protect bacterial substrates in order to benefit SPO1 development. The fact that the ClpCP protease and its adaptor proteins are involved in both regulatory (e.g. transcription factors) and general (misfolded damaged proteins) proteolysis (28) would support the view that a competing ‘xenogeneic’ adaptor protein such a Gp53 would have detrimental pleiotropic effects on the growth of *B. subtilis* cells (Fig. 1).

The ClpC and ClpP proteins of Gram-negative and Gram-positive bacteria have recently been recognised as viable targets for antibiotic discovery and a number of naturally-occurring antibacterial products deregulate the respective activities of ClpC or ClpP resulting in bacterial cell death (29, 30). With the emerging interest in the use of phages and phage encoded proteins as source of alternatives to antibiotics, this study reveals that the ClpCP protease of *B. subtilis* and homologs in other bacteria can be subjected to xenogeneic dysregulation by phage derived factors and adds Gp53 to the growing list of naturally-occurring antibacterial products that target the bacterial protein degradation machinery.

## Materials and Methods

### Plasmids, strains and proteins

All the plasmids used in this study for protein expression and the BTH assays were generated using standard molecular biology procedures and are detailed in Table S1. The pSCBAD-Gp53 was made by Gibson assembly (31): The pSC101 plasmid (32) was modified by inserting the regulatory region of pBAD33 (*araC* promoter region, multiple cloning site and the rrnB T2 terminator) between restriction sites *Xho*I and *Nsi*I. All proteins used in this study were purified by either Ni-affinity chromatography (for 6His tagged proteins i.e. Gp53, MecA, and ClpP) or anti-FLAG M2 affinity resin (for FLAG tagged proteins i.e. ClpC) using standard molecular biology procedures. The details of plasmids used for protein purification are shown in Table S1. All the strains used in this study are shown in Table S2.

### Bacterial growth assays

Unless otherwise stated, *B. subtilis* cultures were grown in 2xYT medium (Sigma) with 2% (w/v) glucose and appropriate antibiotics at 37 °C. For the experiments shown in Fig. 1 and Fig. 2*A*, seed cultures were grown at 37 °C, shaking at 700 rpm for 16-18 hours in a THERMOstar (BMG Labtech) plate incubator by directly inoculating a colony into 200 µl of 2xYT medium containing 5 µg/ml chloramphenicol and 2% (w/v) glucose (to prevent leaky expression from pHT01 vector) into a 48-well plate (Greiner). The growth curves were also performed in 48-well plates in a SPECTROstar Nano Absorbance multiwell plate reader (BMG Labtech): the seed cultures were OD_600_-corrected to 0.025 in 200 µl of fresh 2xYT medium containing 5 µg/ml chloramphenicol, 2% (w/v) glucose, and either water or 1 mM IPTG to induce the expression of SPO1 host takeover genes. Cultures were incubated at 37°C, shaking at 700 rpm. At least three biological and technical replicates were performed.

### Pull down assays

These were performed as previously described by (24) using proteins specified in the main text and figures, with the following amendments: binding buffer (25 mM NaH2PO4, 50 mM NaCl, 5 mM imidazole, 5% glycerol at pH 7), wash buffer (25 mM NaH2PO4, 50 mM NaCl, 15 mM imidazole, 5% glycerol at pH 7) and samples were eluted by adding 50 µl of Laemmli 2x concentrate SDS Sample Buffer to beads and boiled for 5 minutes prior to analysis by SDS-PAGE.

### Bacterial two-hybrid interaction assays

These were carried out using the Bacterial Adenylate Cyclase-based Two-Hybrid (BACTH) system (Euromedex) and were conducted as per manufacturer’s guidelines. Briefly, recombinant plasmids encoding proteins of interest fused to the T25 or T18 domain of adenylate cyclase were transformed into competent DHM1 cells (see Table S1 for details of plasmids used). Transformants were grown in a 96-well plate in LB medium containing ampicillin (100 µg/ml), kanamycin (50 µg/ml), and IPTG (0.5 mM), overnight at 30 °C. Each culture was then diluted 1:5 in Z buffer (45 mM Na2HPO4-NaH2PO4 pH 7, 10 mM KCl, 2 mM MgSO4.7H2O, 40 mM β-mercaptoethanol) and cells were permeabilised using 0.01% (w/v) SDS and 10% (v/v) chloroform. Each culture was again diluted 1:4 in Z buffer and equilibrated at 28 °C, before adding 0.4% (v/v) o-nitrophenol-β-galactoside (ONPG). Reactions were carried out in a SPECTROstar Nano Absorbance multiwell plate reader (BMG Labtech) at 28 °C for 20 minutes, with measurement of OD420nm every 1 minute. The β-galactosidase activity is given in Miller units, with one Miller unit corresponding to 1 nM ONPG hydrolysed per minute at 28°C (after accounting for OD_600_ correction and dilution factors). At least three biological and technical replicates were performed for each measurement.

### ATPase assays

The ATPase assay is based on colorimetric measurement of the concentration of inorganic phosphate (Pi) from the hydrolysis of ATP. Reactions were carried out at 37°C for the specified times in buffer containing 100 mM KCl, 25 mM Tris-HCl at pH 8.0, 5 mM MgCl2, 0.5 mM DTT, 0.1 mM EDTA, 0.5 µg/µl BSA and 4 mM ATP; ClpC, MecA, and/or Gp53 were added at concentrations indicated in figures and figure legends. The amount of Pi in the reaction was then quantified using PiColorLock^TM^ detection reagent (Innova Biosciences) as per manufacturer’s guidelines. The data were corrected for buffer only values to account for any spontaneous degradation of ATP. At least three biological and technical replicates were performed for each reaction.

### ClpCP mediated protein degradation assays

These were conducted exactly as described previously in (15). The protein components were present at the amounts indicated in the figure legends.

### SPO1 infection assays

Seed cultures of bacteria were grown at 37 °C, shaking at 700 rpm for 16-18 hours in a THERMOstar (BMG Labtech) plate incubator by directly inoculating a colony into 1 ml of 2xYT medium into a 24-well plate (Greiner). The infection curves were also performed in 24-well plates in a SPECTROstar Nano Absorbance multiwell plate reader (BMG Labtech). The seed cultures were OD_600_-corrected to 0.05 in 1 ml of fresh 2xYT medium and incubated at 37 °C shaking at 700 rpm. At OD_600_ 0.2 SPO1 lysate was added in a 1:1 ratio of bacterial cells: phage particles and OD_600_ measurements taken every 10 minutes until full lysis of the bacterial culture occurred. At least three biological and technical replicates were performed.

## Acknowledgments

We thank Charles Stewart for providing us the SPO1 phage. This work was supported by Wellcome Trust Investigator Award 100958 to S.W. and a Medical Research Council Ph.D. studentship to N.M. Work in the laboratory of K.T. was supported by the Deutsche Forschungsgemeinschaft (Tu106/6-2, Tu106/8-1) and I.H. received a PhD fellowship of the Hannover School for Biomolecular Drug Research (HSBDR).

## Author contributions

N.M., I.H., K.T. and S.W. designed research; N.M., I.H., L.B. and S.N. performed research; N.M., I.H., K.T. and S.W. analysed data and N.M., I.H., K.T. and S.W. wrote the paper.

## Supplementary figure legends

**Fig. S1.**
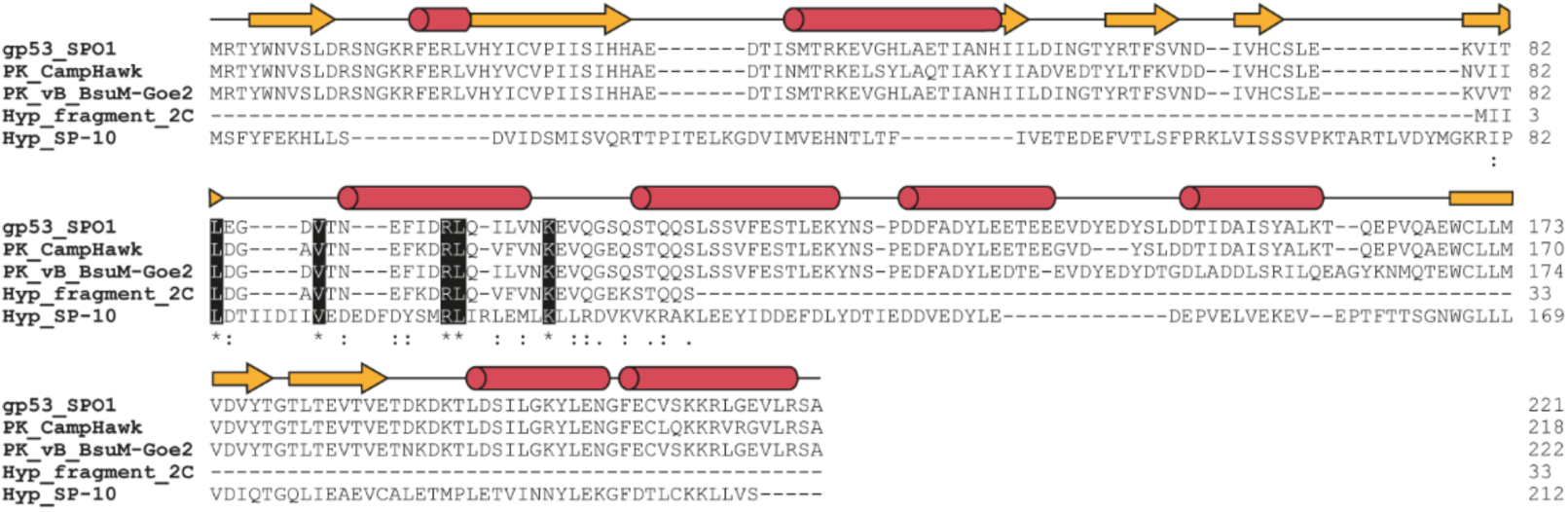
Alignment of amino acid sequences from Gp53-like proteins from *B. subtilis* phages. The localization of the β-strands and α-helices in Gp53 are indicated by yellow arrows and red cylinders, respectively. Residues which are conserved across all sequences that were targeted for mutagenesis are highlighted in black.

**Supplementary Table 1.**
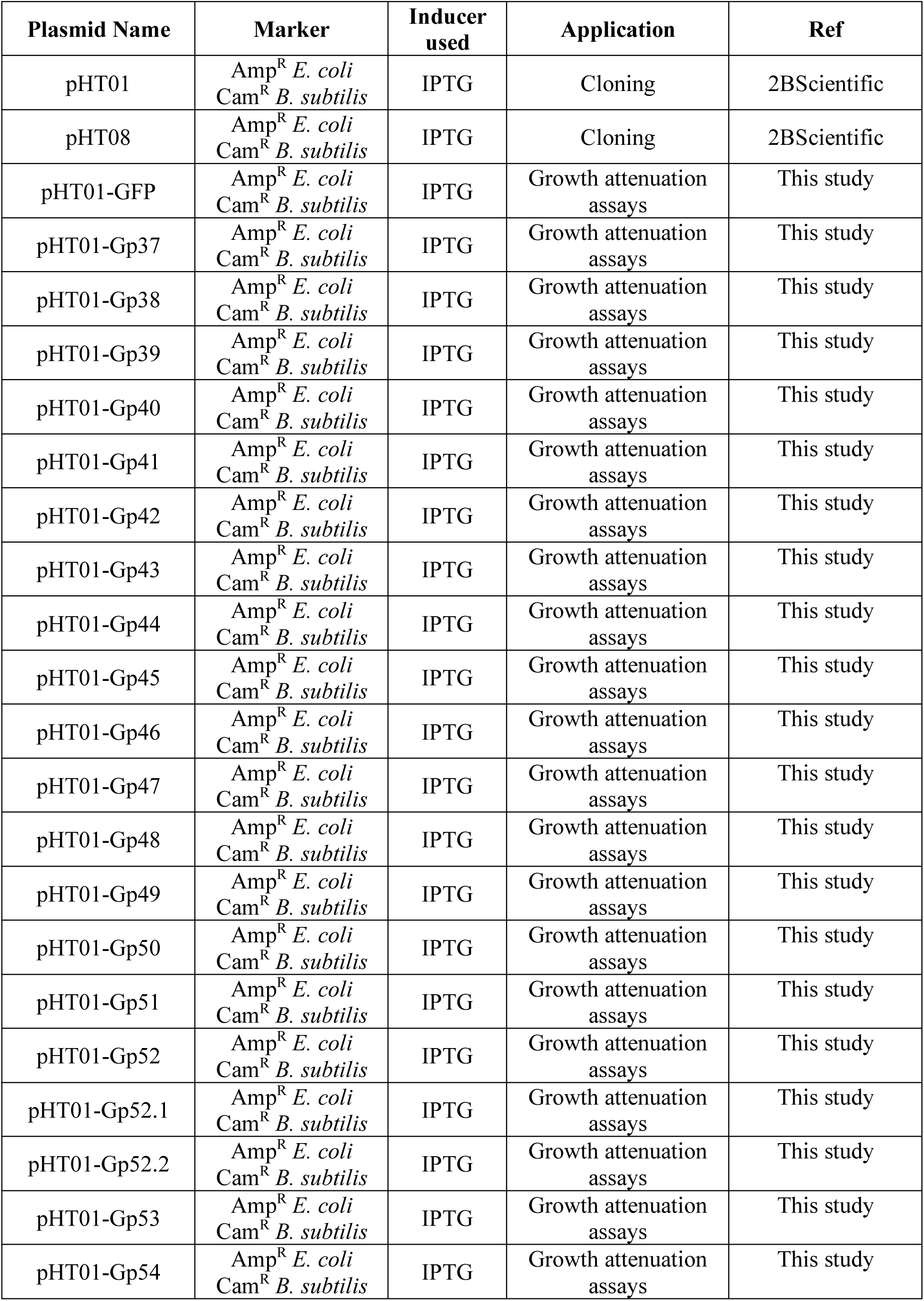
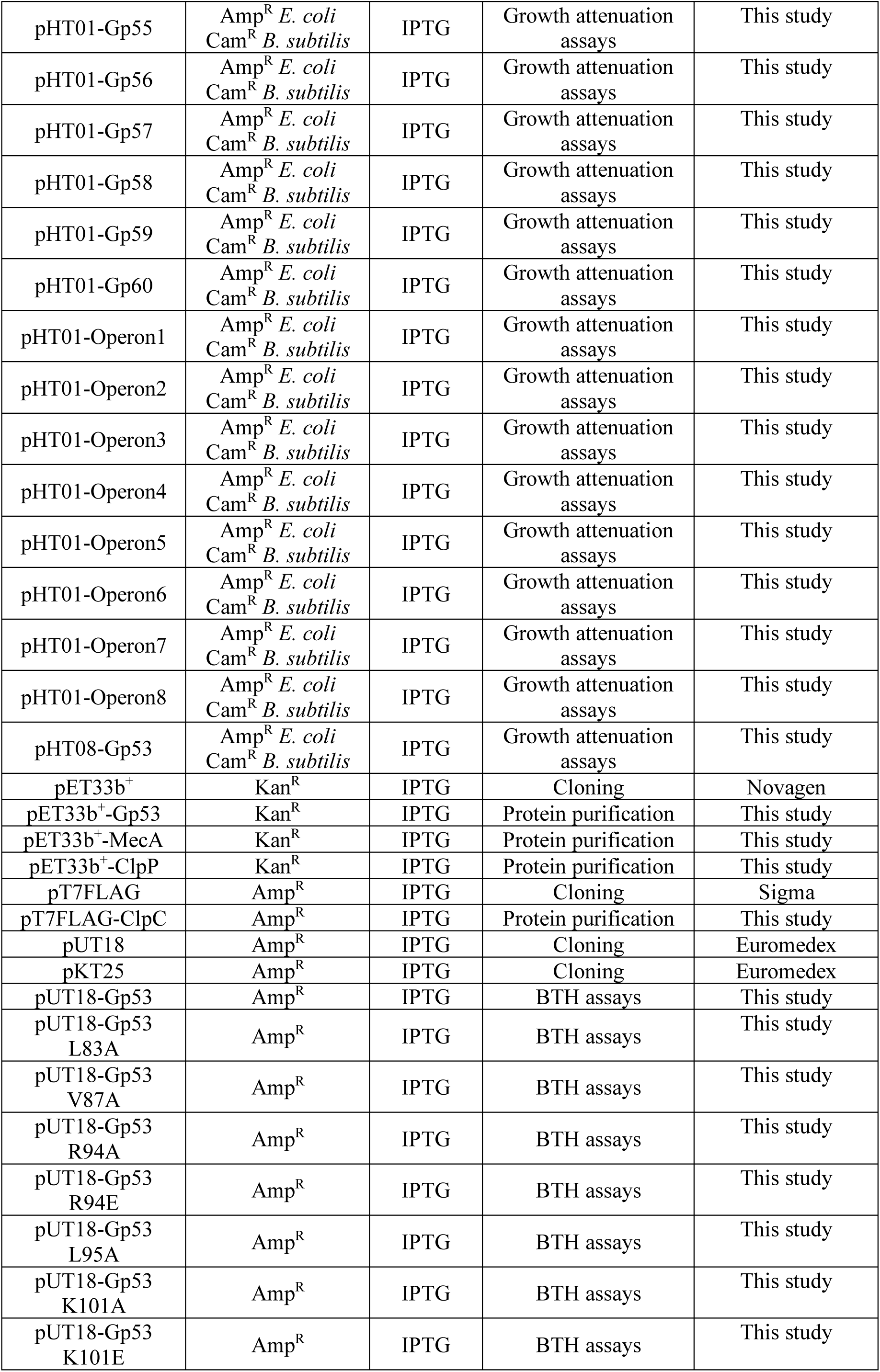
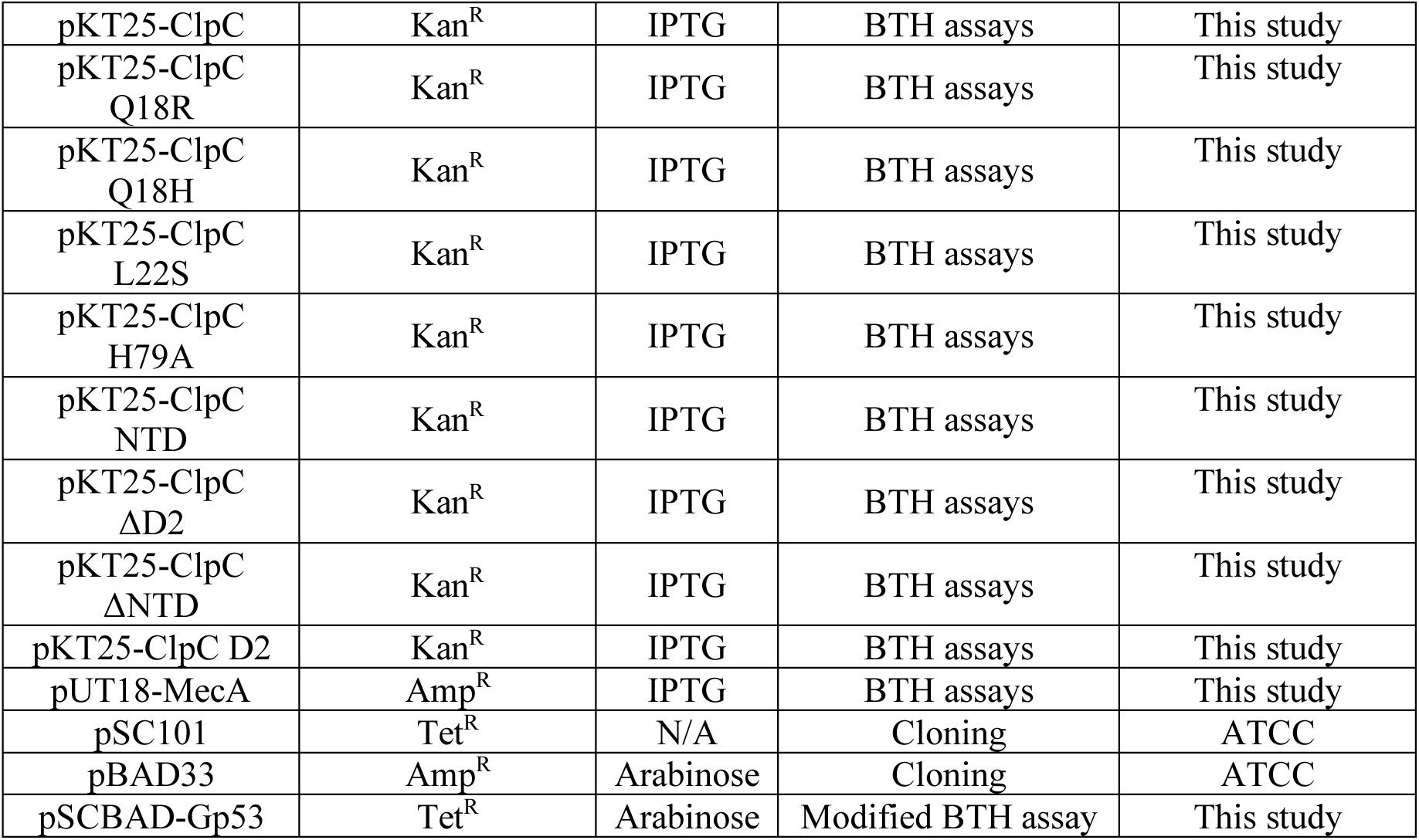
Plasmids used during this study

**Supplementary Table 2.**
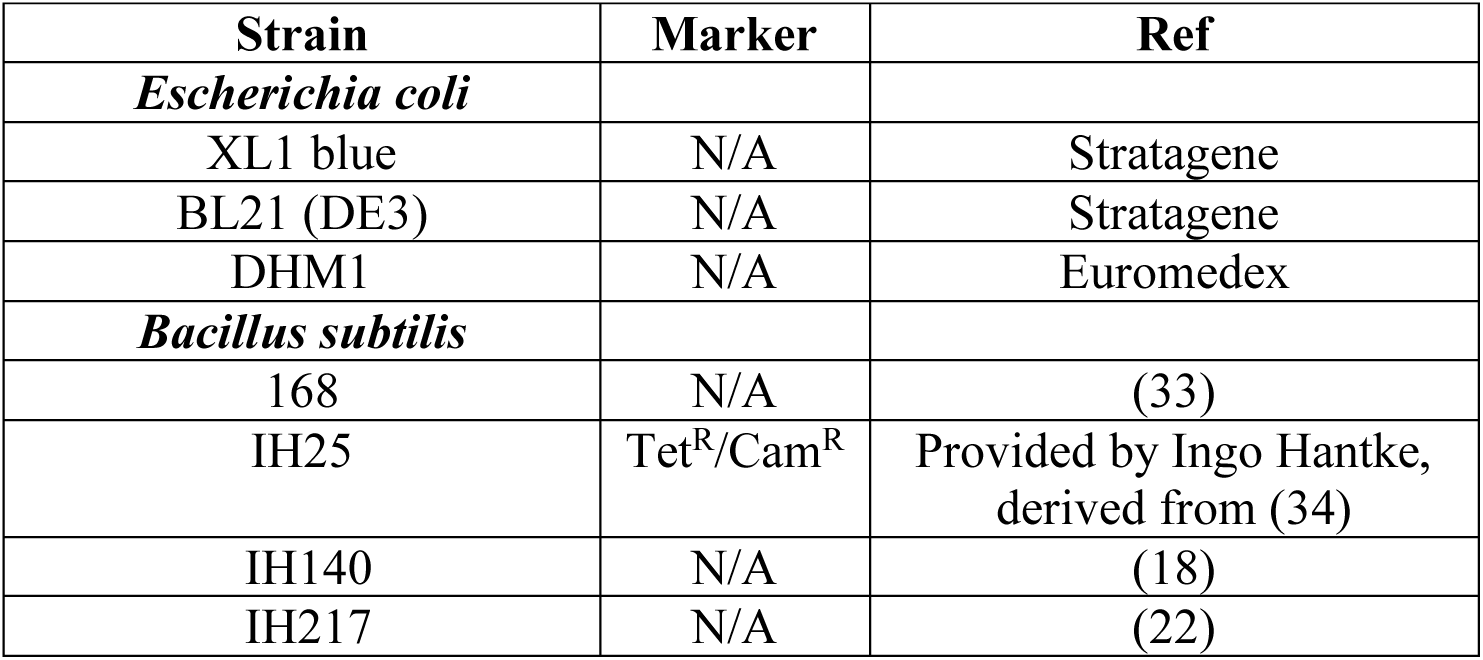
Strains used in this study.

